# Direct construction of sparse suffix arrays with Libsais

**DOI:** 10.1101/2025.02.24.639849

**Authors:** Simon Van de Vyver, Tibo Vande Moortele, Peter Dawyndt, Bart Mesuere, Pieter Verschaffelt

## Abstract

Pattern matching is a fundamental challenge in bioinformatics, especially in the fields of genomics, transcriptomics and proteomics. Efficient indexing structures, such as suffix arrays, are critical for searching large datasets. While sparse suffix arrays offer significant memory savings compared to full suffix arrays, they typically still require the construction of a full suffix array prior to a sampling step, resulting in substantial memory overhead during the construction phase. We present an alternative method to directly construct the sparse suffix array using a simple, yet powerful text encoding, in combination with the widely used Libsais library. This approach bypasses the need for constructing a full suffix array, reducing memory usage by 63% and construction time by 55% when building a sparse suffix array with sparseness factor 3 for the entire UniProt knowledgebase. The method is particularly effective for applications with small alphabets, such as a nucleotide or amino acid alphabet. An open-source implementation of this method is available on GitHub, enabling easy adoption for large-scale bioinformatics applications.

## Introduction

Efficiently matching large amounts of short sequences against large reference databases is a fundamental challenge in bioinformatics, particularly in the fields of genomics, transcriptomics, and proteomics. As datasets grow in size and complexity, this challenge becomes increasingly important. Over the years, a lot of specialized algorithms, such as the Knuth-Morris-Pratt [1] (KMP) and Boyer–Moore–Horspool [2] algorithms, have been developed. These algorithms match sequences (consisting of *m* characters) in a text (consisting of *n* characters) in a worst case time complexity of *O*(*n* + *m*) and *O*(*n m*) respectively. However, such approaches require traversing the entire text to locate all matches, which becomes computationally expensive when the text is large (*n* >> *m*). To address this, index structures, such as suffix trees [3] and suffix arrays [4], have been introduced. These index structures preprocess the text, allowing short sequences to be matched in *O*(*m*) and *O*(*m* log(*n*)) time respectively.

A suffix tree is a compressed trie that stores all suffixes of a given text, allowing patterns to be located by descending the tree from the root along the path matching the pattern. Despite their utility, suffix trees often have a memory footprint that is several times the size of the original dataset, making them less suitable for large bioinformatics datasets. Suffix arrays provide a space-efficient alternative by storing a sorted list of all text suffixes, instead of directly modelling the tree structure. A binary search over the suffix array allows rapid identification of the range of suffixes starting with the desired sequence. Although suffix arrays are more memory-efficient than suffix trees, their memory requirements can still be the limiting factor for large datasets. For example, a suffix array constructed for the UniProt knowledgebase [5] version 2024.04, where all protein sequences are concatenated and separated by a unique character, requires approximately 650 GB of memory, despite the total dataset being only around 82 GB. Sparse suffix arrays can be used to reduce this memory footprint even further.

A sparse suffix array (SSA) retains only suffixes at every *k*-th position in the text, where *k* is the sparseness factor. This means that an SSA of a text of length *n* contains only 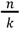 entries instead of *n*, reducing space requirements by a factor of *k*. However, performing lookups in a SSA is slower than in a full SA because additional steps are required to simulate the full SA. The parameter *k* represents a trade-off, balancing reduced memory usage against slower search speed.

Several highly optimized software libraries, such as Libdivsufsort [6] and Libsais [7], are available for the construction of suffix arrays. While very popular, these libraries do not natively support the construction of sparse suffix arrays. The traditional approach to constructing a sparse suffix array involves first constructing the full suffix array and then sampling it. This sampling step selects only every *k*-th entry from the full suffix array to create the sparse suffix array while discarding the rest. Because the full suffix array must still be built beforehand, this method does not reduce peak memory usage, which remains as high as that of constructing the full suffix array. In this article, we introduce an alternative approach for the direct construction of sparse suffix arrays. This method provides fast sparse suffix array construction across various domains, including genomics, transcriptomics, and proteomics. This method reduces memory usage by 63% and execution time by 55% for the SSA construction for the UniProtKB with sparseness factor 3.

## Methods

To optimize memory usage and accelerate the construction of sparse suffix arrays with sparseness factor *k*, we first apply a transformation to the input text. This transformation maps each non-overlapping k-mer to a unique integer, ensuring that lexicographically smaller groups correspond to smaller integer values and preserving equivalence. As a result, the effective length of the transformed text is reduced by a factor *k*. One approach to such a transformation is to first assign a unique unsigned integer to each character in the alphabet, using the minimal number of bits necessary for representation. Next, each k-mer is bit-packed [8] into a single unsigned integer, using the smallest possible data size required to represent the group. The bit-packing process is structured such that the leftmost character in each group occupies the most significant bits of the integer, with each subsequent character assigned to the next most significant bits still available. This ensures that lexicographic order is preserved, with smaller groups consistently corresponding to smaller integer values. For the canonical amino acid alphabet consisting of 20 different characters, where each character requires 5 bits, grouping *k* = 3 characters together results in 15 bits per group. These 15 bits can be efficiently stored in a 16-bit unsigned integer within the transformed text. Consequently, the length of the transformed text is reduced by a factor of *k* = 3.

We then build the suffix array for this sequence of encoded unsigned integers. The result is a SSA of the original input text, equivalent to a sampled SSA derived from a full suffix array. However, it is constructed directly, eliminating the need to first build the full suffix array (see Figure 1).

**Figure 1:**
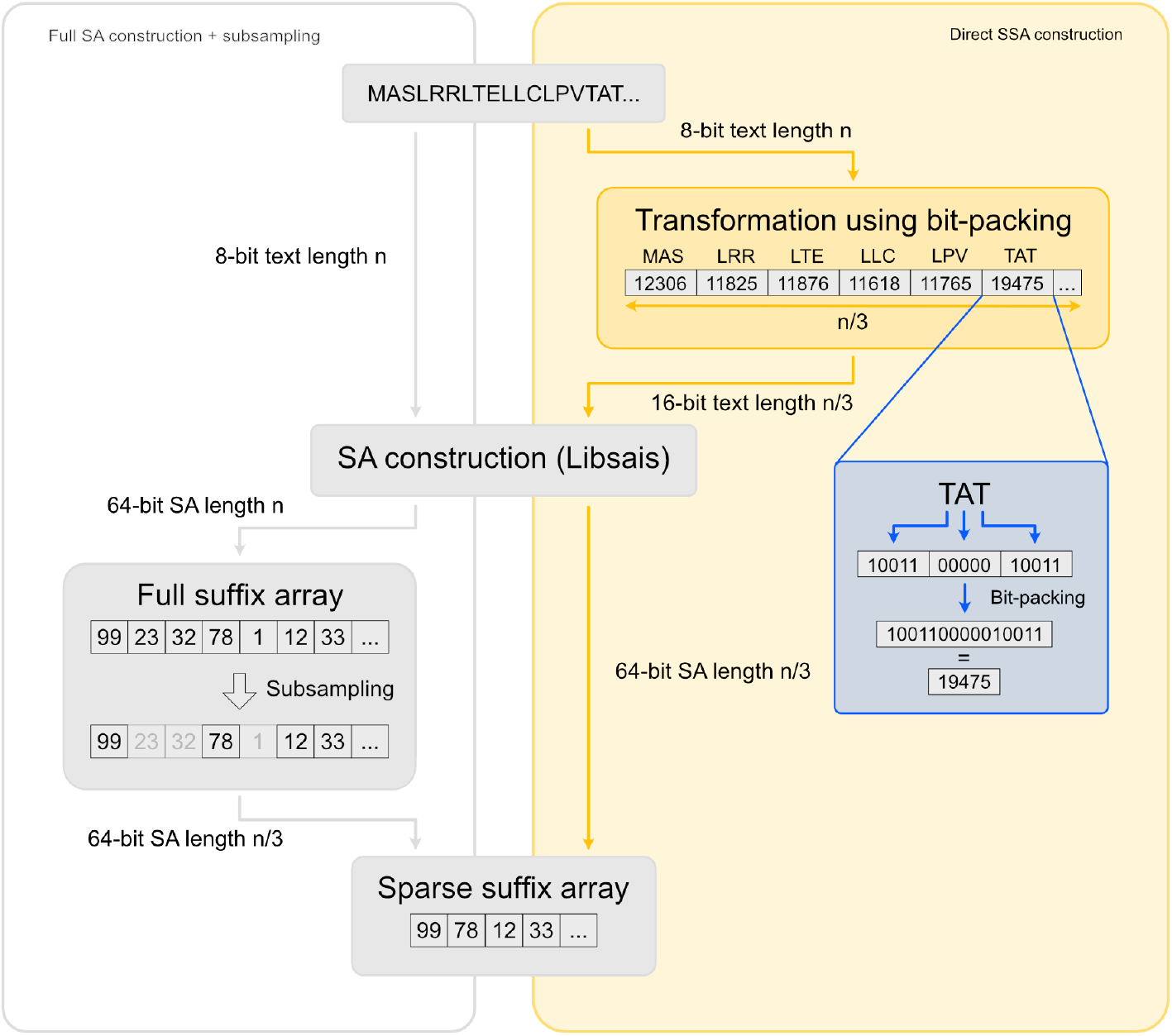
Overview of the standard method (left) using subsampling and the direct method (right) using a text transformation to directly construct the sparse suffix array for sparseness factor 3. The text transformation reduces the length of the text from n 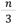.

This method for direct construction of a sparse suffix array can be used for all suffix array construction libraries, but its efficiency depends on the specific implementation. We use Libsais due to its speed and memory efficiency, and the following complexity analysis is based on its internal algorithm, induced suffix sorting. With Libsais, this approach changes the memory complexity from 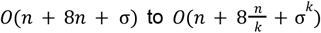,where *n* is the length of the original text and σ is the size of the alphabet. The first and second term in each complexity expression correspond to the size of the input text and the SSA respectively. The last term corresponds to the memory required for bucket allocation performed as part of the induced sorting algorithm, which is the algorithm used internally by Libsais. These buckets store occurrences of each character in the transformed alphabet, where the transformation groups k-mers into a single unit. As a result, the number of distinct characters in the transformed alphabet increases from σ to σ^*k*^, leading to a change in bucket memory requirements.

For small values of the sparseness factor *k* and small alphabet sizes, this approach yields substantial reductions in both memory usage and execution time, as the effective input length for suffix array construction decreases. However, if *k* or σ is too high, the memory savings may diminish or even increase due to the allocated buckets. In these cases, it is advantageous to lower the sparseness factor during construction to a divisor of *k* and apply sampling post-construction to achieve the desired sparseness. This flexible approach allows for efficient suffix array construction across a variety of sparseness factors and alphabet sizes.

## Results

We implemented this method in C to ensure efficient performance and seamless integration into existing bioinformatics workflows. Libsais-packed^1^ [9], our open-source implementation of the direct construction sparse suffix arrays demonstrates a large performance boost compared to the traditional method of SSA construction. This approach is particularly effective for datasets with small alphabets, such as genomic, transcriptomic and proteomic sequences. To evaluate its impact, we performed benchmarking tests using two datasets: UniProtKB [5] 2024.04 and a human reference genome.

The UniProtKB database consists of TrEMBL and SwissProt, containing approximately 245 million proteins. All protein sequences are concatenated using a separator character, resulting in a total text length of approximately 82 GB. The reference genome (genome assembly GCF_000001405.40; GRCh38.p14) was downloaded from NCBI and contains a full sequence of approximately 3 GB. To focus on the core nucleotide data, all ‘N’ characters were removed, keeping only the nucleobases (A, C, G, and T).

### Benchmark Setup

The UniProtKB dataset was prepared by concatenating all proteins using a dash (‘-’) as a separation character. For the reference genome, we used a sequence where all N’s were removed, resulting in a simplified sequence containing only A, C, G, and T, which can be encoded in the transformed text using only 2 bits per character. The sparse suffix array construction was evaluated with varying sparseness factors, comparing the standard method using subsampling with the new method using the bit-packing text transformation technique.

### UniProtKB Dataset Performance

Figure 2a and 2b illustrate the execution time (a) and memory usage (b) for sparse suffix array construction on the UniProtKB dataset. Direct construction with Libsais-packed substantially reduces memory usage, with a peak reduction of 73% at sparseness factor 5 compared to the standard approach. As the sparseness factor increases, memory consumption drops sharply at first but starts rising again at sparseness factor 6, reflecting the growing overhead of handling an expanded alphabet in the transformed text. Execution time follows a similar trend, with the method achieving its peak reduction of 60% at sparseness factor 4. Beyond this point, the gains diminish as the alphabet size increases, introducing additional processing overhead. These results demonstrate that increasing the sparseness factor improves efficiency. However, once the alphabet size expands too much, it introduces overhead, reducing the memory and time savings.

**Figure 2:**
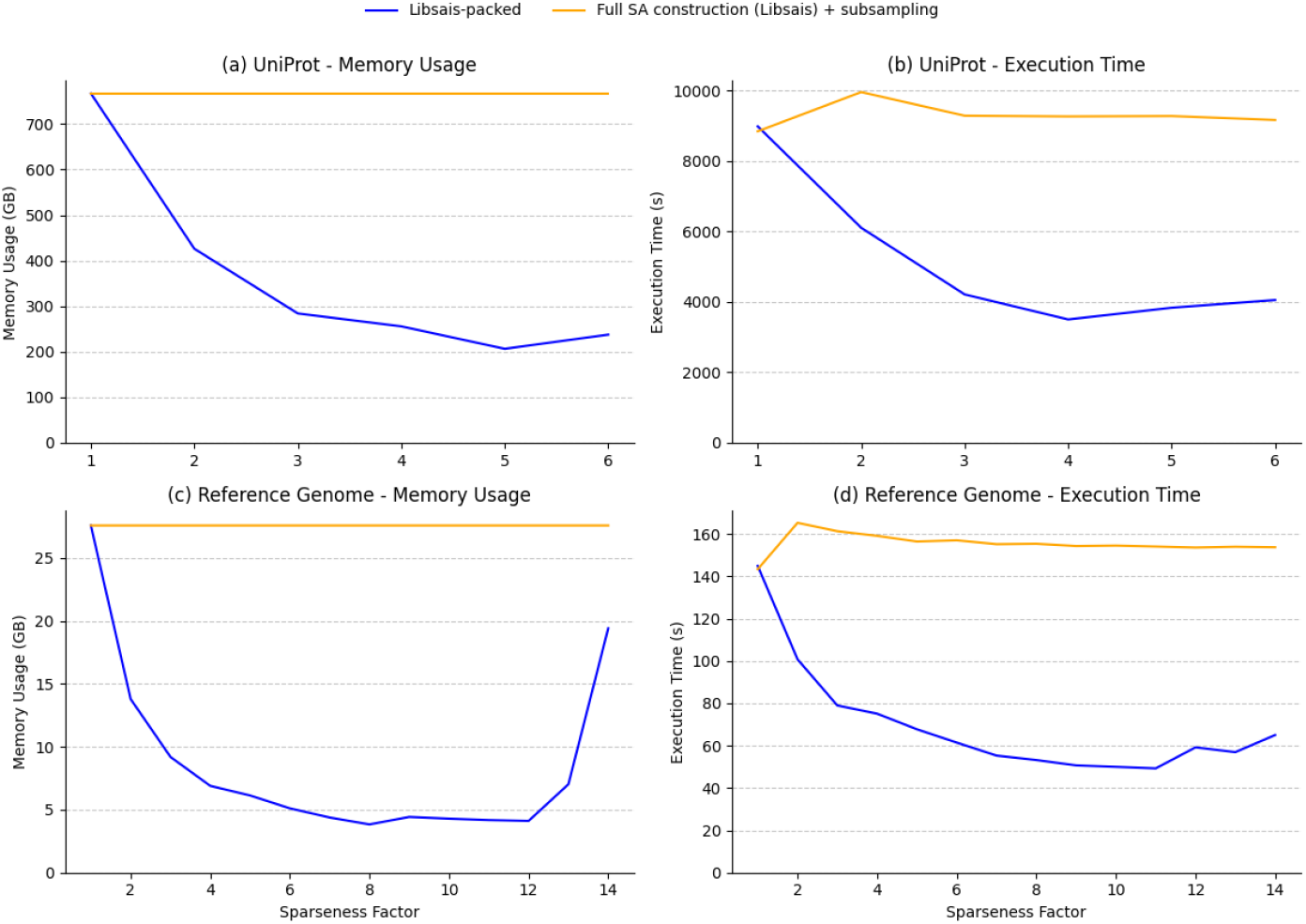
Memory usage (a) and execution time (b) of the SSA construction for the Uniprot dataset, and memory usage (c) and execution time (d) of the SSA construction for a reference genome for varying sparseness factors. The orange lines show the performance of the traditional SSA construction, using full SA construction with Libsais and subsampling, The blue lines show the performance of Libsais-packed, which constructs the SSA directly through a text transformation.

### Reference Genome Performance

Figure 2c and 2d show the execution time (c) and memory usage (d) for sparse suffix array construction of the reference genome under the same conditions as for the UniProtKB database. The overall trends mirror those observed for the UniProtKB dataset. Memory usage initially decreases steeply for direct construction, reaching its lowest point at sparseness factor 8, where it is reduced by 86%. Beyond this point, the growing alphabet size leads to increased memory requirements. Similarly, execution time improves as the sparseness factor increases, with a peak reduction of 68% at factor 11, before the overhead from the expanded alphabet begins to counteract the gains.

## Conclusion

In this article, we introduced an efficient method for the construction of sparse suffix arrays for large datasets. Central to this approach is the introduction of a simple text transformation, which encodes k characters of the input text into a compact representation, which then serves as input to Libsais. This method reduces the length of both the input text and the resulting suffix array by a factor of k. Libsais-packed [9] is a C implementation of this method that can be used as a drop-in replacement for Libsais.

Libsais-packed delivers significant improvements in both memory usage and execution time for the construction of a sparse suffix array, as long as the alphabet size remains manageable, such as in genomics and proteomics. The benefits diminish when the alphabet size becomes excessively large, emphasizing the need to carefully select parameters for the specific dataset.

## Footnotes

1 Available at https://github.com/unipept/libsais-packed

## References

[1] D. E. Knuth, J. H. Morris, Jr., and V. R. Pratt, “Fast Pattern Matching in Strings,” SIAM J. Comput., vol. 6, no. 2, pp. 323–350, Jun. 1977, doi: 10.1137/0206024.

[2] R. N. Horspool, “Practical fast searching in strings,” Softw. Pract. Exp., vol. 10, no. 6, pp. 501–506, 1980, doi: 10.1002/spe.4380100608.

[3] P. Weiner, “Linear pattern matching algorithms,” in 14th Annual Symposium on Switching and Automata Theory (swat 1973), Oct. 1973, pp. 1–11. doi: 10.1109/SWAT.1973.13.

[4] U. Manber and G. Myers, “Suffix Arrays: A New Method for On-Line String Searches,” SIAM J. Comput., vol. 22, no. 5, pp. 935–948, Oct. 1993, doi: 10.1137/0222058.

[5] The UniProt Consortium, “UniProt: the Universal Protein Knowledgebase in 2025,” Nucleic Acids Res., vol. 53, no. D1, pp. D609–D617, Jan. 2025, doi: 10.1093/nar/gkae1010.

[6] Y. Mori, Libdivsufsort. [Online]. Available: https://github.com/y-256/libdivsufsort

[7] I. Grebnov, Libsais. [Online]. Available: https://github.com/IlyaGrebnov/libsais

[8] D. Salomon and G. Motta, Handbook of Data Compression. Springer Science & Business Media, 2010.

[9] unipept/libsais-packed. (Feb. 19, 2025). C. Unipept. Accessed: Feb. 19, 2025. [Online]. Available: https://github.com/unipept/libsais-packed

